# Minor differences in peptide presentation between chicken MHC class I molecules can explain differences in disease susceptibility

**DOI:** 10.1101/2022.03.11.484051

**Authors:** Lingxia Han, Shaolian Wu, Weiyu Peng, Min Zhao, Can Yue, Wanxin Wen, Wenbo Cai, Ting Zhang, Hans-Joachim Wallny, David W. Avila, William Mwangi, Venugopal Nair, Nicola Ternette, Yaxin Guo, Yingze Zhao, Yan Chai, Jianxun Qi, Hao Liang, George F. Gao, Jim Kaufman, William J. Liu

## Abstract

The highly polymorphic classical major histocompatibility complexes (MHCs) can confer resistance or susceptibility to diseases. The chicken MHC is known to confer decisive resistance or susceptibility to various economically-important pathogens, including the iconic oncogenic herpesvirus that causes Marek’s disease (MD). Only one classical class I gene, BF2, is expressed at a high level in chickens, so it was relatively easy to discern a hierarchy from well-expressed thermostable fastidious specialist alleles to promiscuous generalist alleles that are less stable and expressed less on the cell surface. The BF2*1901 is more highly expressed and more thermostable than the closely-related BF2*1501, but the data for peptide repertoire available did not obviously correlate as expected. Here, we confirm for newly-developed chicken lines that the chicken haplotype B15 confers resistance to MD compared to B19. Using gas phase sequencing of peptide pools, and using immunopeptidomics involving mass spectroscopy, we find that the BF2*1901 binds a greater variety of amino acids in some anchor positions than BF2*1501. However, by X-ray crystallography, we find that the peptide-binding groove of BF2*1901 is narrower and shallower. Though the self-peptides bound to BF2*1901 may appear more various than those of BF2*1501, the structures show that the wider and deeper peptide-binding groove of BF2*1501 allows it to tightly accept many more peptides overall, correlating with the expected hierarchies for expression level, thermostability and MD resistance. Moreover, the α2 helix of BF2*1901 is higher than BF2*1501, potentially reducing the number of T cell clones that can recognize this fastidious class I molecule.

**IMPORTANCE:** Disease susceptibility mechanism is complicated, but chicken infection of Marek’s disease (MD) is one of ideal models, considering the only one highly expressed classical MHC class I, BF2. The different susceptibility of the two close alleles BF2*1901 and BF2*1501 with minor difference of expression and thermostability is still unfathomed. Here, we confirm B15 chicken confers resistance to MD compared to B19. But the BF2*1901 binds a broader variety of anchoring peptides than BF2*1501. This mystery was solved by the structural determination of the two molecules with one similar peptide. The wider and deeper peptide-binding groove of BF2*1501 allows it to tightly accept many more peptides overall, which is concordant to the expected hierarchies for expression level, thermostability and MD resistance.

## INTRODUCTION

The global pandemic of COVID-19 among humans caused by the coronavirus SARS-CoV-2 has emphasized the importance of understanding the mechanisms of resistance against viral pathogens(1). Compared to roughly 7 billion humans, there are estimated to be over 80 billion chickens alive each year, most of which are potentially subject to local epidemics by a variety of economically-important viral diseases (http://www.fao.org/faostat/en/#data/QL). The first coronavirus ever described causes infectious bronchitis in chickens and is still a major problem for commercial flocks(2, 3), but the iconic chicken pathogen is Marek’s disease virus (MDV), an oncogenic herpesvirus for which most commercial chickens are vaccinated and which still causes major economic losses due to changes in virulence and tropism (4–6). Much ongoing research is dedicated to determining the genetic loci responsible for resistance to Marek’s disease (7–9), but the BF-BL region within the B locus, which is clearly the functional equivalent of the major histocompatibility complex (MHC), has been known for decades to determine resistance and susceptibility (10–12).

In contrast to humans and other typical mammals, the chicken MHC is small and simple, and can determine striking resistance or susceptibility to various economically-important infectious diseases (13). Compared to humans which express multigene families of classical MHC class I and class II molecules, there are only two classical class I (BF) genes and two classical (BL) class II genes, and only one of each is expressed at a high level. The BF2 molecule is well-expressed and is the major ligand for cytotoxic T lymphocytes, while the BF1 molecule acts as a ligand for natural killer (NK) cells and is relatively poorly expressed if at all(14). The presence of dominantly-expressed classical MHC molecules whose properties can determine the immune response has been suggested to be one reason why the chicken MHC has such strong genetic associations with infectious diseases(15), although other closely-linked genes may also be involved(16, 17).

The simplicity of the chicken MHC compared to typical mammals has allowed the discovery of some fundamental properties of classical MHC molecules, in particular the properties of class I molecules leading to the proposal of generalist and specialist alleles (18–20). In chickens, there is a clear hierarchy of class I alleles from so-called fastidious molecules that bind a narrow range of peptides, are relatively stable with the peptides naturally bound and have a relatively high cell surface expression compared to so-called promiscuous molecules that bind a wider variety of peptides, are overall less stable and have a lower expression at the cell surface (15, 18, 21–23). It has been relatively easy to understand the size of the peptide repertoire from the structures of the chicken class I molecules(15, 18, 22, 24–26), although peptide transport by TAP molecules and peptide editing by tapasin (TAPBP) may also contribute(23, 27, 28). The chicken MHC haplotypes with promiscuous class I alleles are generally associated with resistance to a variety of infectious viruses, including those responsible for Marek’s disease, infectious bronchitis, avian influenza and Rous sarcoma (19, 20)

A similar hierarchy of human classical class I alleles has been discerned (18-20, 29-31). For human class I alleles, the original observation was that fastidious class I molecules (so-called elite controller alleles) correlated with slow progression from HIV infectious to AIDS (19, 20, 29, 30), apparently due to binding special pathogen peptides that the virus cannot change for immune evasion without lowering viral fitness (32, 33). Based on assays of tapasin-dependence (34), these protective human alleles were correlated with dependence on the class I-bespoke chaperone tapasin (or TAPBP) in the peptide-loading complex (PLC)(18, 19, 29, 35). The results in chickens and humans led to the concept of generalist class I alleles that generally protect from many viral pathogens by binding a wide variety of peptides and specialist alleles that protect from particular pathogens by binding special peptides (18–20, 29). Most recently, it was found that promiscuous class I alleles in humans correlate with slow progression to AIDS if the elite controller alleles are removed from the analysis (35). The presence of fastidious class I alleles in chickens may also be explained by resistance to particular pathogens, although the evidence is not definitive(20).

That chicken MHC haplotypes are in a hierarchy with respect to resistance to Marek’s disease is not in question, although the relative placement of particular haplotypes within that hierarchy has been debated (12, 36, 37). It is perhaps a surprise that any consensus could have arisen, given the differences between experiments in the relative virulence of different MHC strains, the route of infection, the measurement of disease, the chicken lines with different genetic backgrounds and the differences even within MHC haplotypes. For example, there are clearly two kinds of B15 haplotypes, those that have a functional BF1 gene and those that do not (15, 22, 38, 39), and the relative resistance to MDV conferred by these haplotypes has not been examined. Moreover, there is evidence the BG1 gene within the chicken MHC can contribute to resistance to virally-induced tumours(16), with many of the infection experiments carried out before the BG1 gene was even discovered (40–42).

There seems to be little disagreement that the B19 haplotype confers the most susceptibility, and most experiments with B15 haplotypes show that it confers susceptibility but less than B19(12, 36, 37). In agreement, the expression level and thermostability are higher for class I molecules on erythrocytes and splenocytes for B19 than B15 (18, 21, 23). However, it has not been obvious from published data on motifs whether BF2 molecules from B19 are more fastidious than B15, nor has there been a structure for the BF2 molecule from a B19 haplotype. Using recently established lines(43), we report the viral levels from a B15 and a B19 chicken line, describe the peptide motifs of the two haplotypes in much more detail than previously, and determine structures for both B15 and B19 with the same peptide as well as with several other peptides, including one B15 structure recently published (24). From these analyses, we confirm that viremia for MDV is higher in the B19 than the B15 chicken lines, and find a structural explanation for greater promiscuity for the BF2 molecule from B15 compared to B19, correlating with the facts that B19 has higher surface expression, greater stability and confers susceptibility to Marek’s disease.

## RESULTS

### The expression of three cytokines is higher in B19 than in B15 naïve SPF chickens

To investigate the transcription level of IFN-γ, IL-18, and IL-10 in the healthy chickens, we developed dqRT-PCR method for the three cytokines; the standard curve showed that the amplification curves in each amplification reaction were standard “S” type, and the amplification efficiency of the target gene and the reference gene was similar, and showed a good linear relationship (S1 Fig.). Based on the quality-controlled dqRT-PCR, mRNA expression of IFN-γ, IL-18, and IL-10, genes were quantified to compare the natural cellular immunological level between SPF B15 and B19 chicken lines. Firstly, the expression levels in PBLs from 14 through 70 d-old chickens showed that B19 chickens transcribe less IFN-γ but more IL-10 than B15 chickens, though the variation was not statistically significant. IL-18 expressed similarly in both the lines except at 56 d-old (S2A-C Fig.).

Secondly, the relative expression levels of IFN-γ, IL-18, and IL-10 in lung and respiratory tract, thymus, bursa of Fabricius, spleen and peripheral blood were detected at 70 d-old, since generally the immune organs of chicken mature at 2 month old. All the three cytokines were expressed in the primary lymphoid organs of thymus and bursal, but only at a low level in respiratory system, spleen and PBLs. The expression levels differed among the tissues and cytokines, with IL-10 and IL-18 transcribed mainly in the bursa and IFN-γ mainly in thymus. Significantly, the three cytokines were expressed more in the corresponding organs of B19 chickens compared to B15 chickens (S3 Fig.).

### Disease susceptibility distinction of B19 and B15 chickens infected by MDV Md5

One-day old chickens were inoculated i.p. with the very virulent Md5 strain of MDV. At 4, 7, and 9 dpi, the number of virus copies within the feather pulps of B19 and B15 chickens retained quite similar. However, at 12 dpi, the virus copy numbers in B19 chickens were much higher than B15 chickens (Fig. 1A). At 20 dpi, 2 chickens died in the B19 group and 1 chicken died in the B15 challenge group. The spleen and kidney of the dead chickens infected with B19 were enlarged, the thymus glands were atrophied, and the liver was congested with the surface color darkened. The livers were atrophied and the kidneys were enlarged in the B15 chickens that died, but the surface color of the livers was lighter.

**Fig 1.**
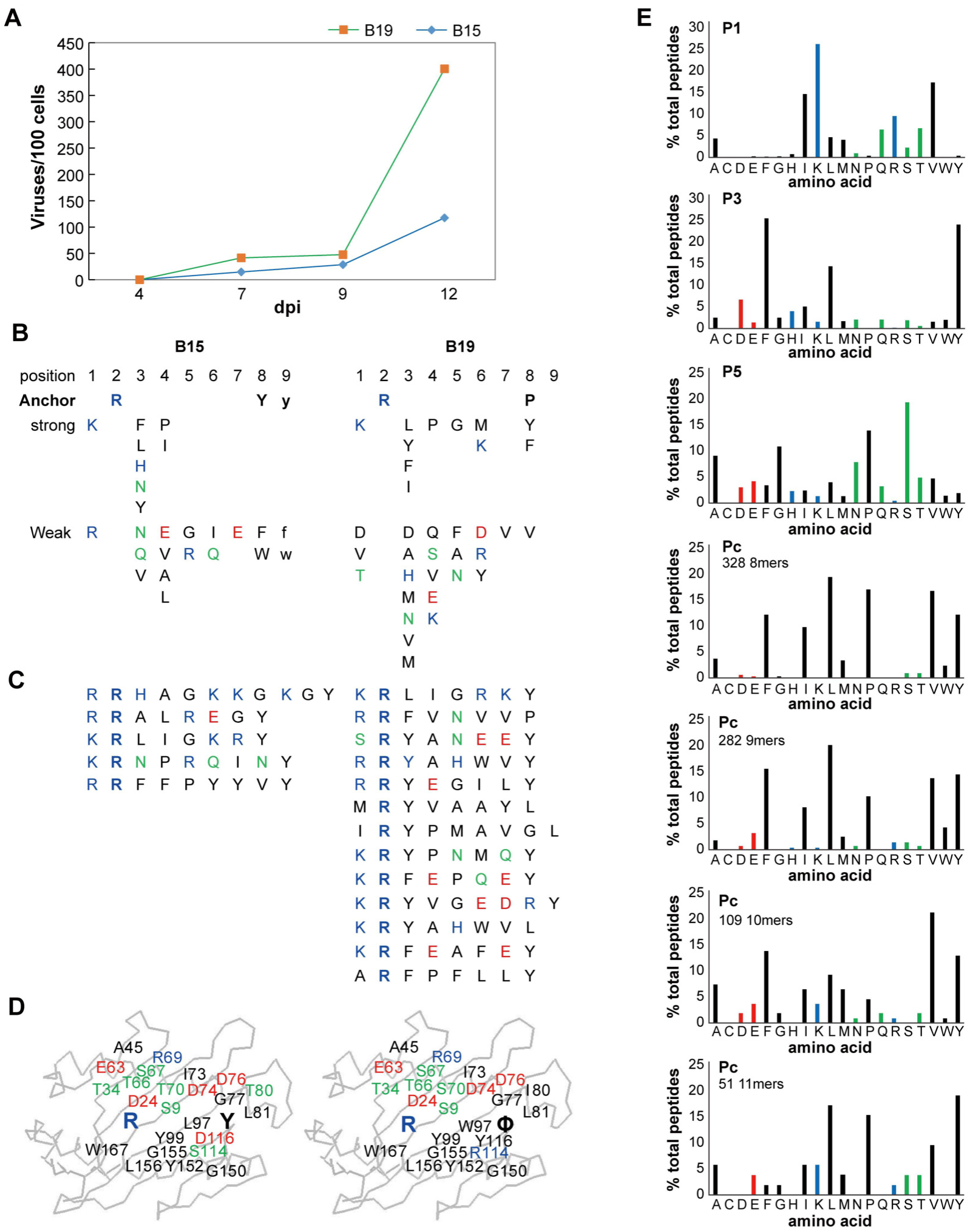
Susceptibility of B15 and B19 SPF chickens to MDV Md5 and peptide preference of BF2*1501 and BF2*1901. A. The dynamic viral load of MDV Md5 in B19 and B15 haplotype chickens at 4, 7, 9, and 12 dpi. B-D. Self-peptides bound to BF2*1501 and BF2*1901 as assessed by gas phase sequencing. B. Sequences of peptides bound to class I molecules isolated from red blood cells determined from peptide pools showing anchor, strong and weak signals. C. Sequences of individual peptides separated by HPLC. D. Peptide anchor residues in large letters superimposed on a wire model of class I α1 and α2 domains with those residues that are both polymorphic and potentially peptide contacts indicated as smaller letters; numbering based on HLA-A2 sequence. Single letter code for amino acids (with Φ for hydrophobic); basic residues in blue, acidic residues in red, polar residues in green, hydrophobic residues in black. Results and analysis for B15 adapted from previous study(15) . E. Analysis of peptides from B19 cells as assessed by immunopeptidomics. Bar graphs showing the frequency (y-axis) of each natural amino acid (single letter code: basic residues in blue, acidic residues in red, polar residues in green, hydrophobic residues in black; x-axis) for peptides eluted from an MDV-transformed cell line MDCC-265L which have Arg at P2 (thus likely to be BF2*1901), for peptide positions P1, P3 and P5, and for the C-terminal amino acid (called Pc or Pω) separated by peptide length and with the number of each length indicated. Original data from experiment described in previous study (18).

### Peptides and peptide motifs from class I molecules of B15 and B19 chickens

As mentioned in the introduction, isolation of class I molecules from chicken blood and spleen cells followed by HPLC and gas phase sequencing of single peptides and peptide pools provided the first glimpses into how chicken class I molecules bind peptides. The first description involved what now might be called fastidious molecules with multiple simple anchor residues, and showed that the class I molecules from the B15 and B19 haplotypes had very similar motifs(21), with an Arg at peptide position 2 (R2) for both, a Tyr at Pc (also called PΩ, in this case P8 or P9) for B15, and a few hydrophobic amino acids (including Leu, Phe, Pro and Tyr) at Pc (P8) for B19. We now know that chickens typically have a BF2 that is the dominantly-expressed class I gene: B19 has a poorly-expressed BF1 gene and most B15 haplotypes have no functional BF1 gene(15, 44), so these gas phase sequencing results reflect the peptides from the BF2 molecule. The B15 peptides and motifs were described in detail (15, 22), but the detailed B19 results are only presented now (Fig. 1B,C). A pool sequence and 13 individual peptides confirm the initial points: both 8mers and 9mers are found for B15, but B19 has mostly 8mers; both B15 and B19 have Arg for P2; B15 is mostly Tyr (with some Phe and other hydrophobic amino acids) at Pc, but B19 has Tyr, Pro, Leu (and some Phe) at Pc. In addition, B15 has entirely basic residues Arg and Lys at P1, while B19 has mostly Lys but some Arg and some hydrophobic residues. Finally, the two motifs fit well with wire models of the class I molecules (as described for B15)(15)(Fig. 1D), predicting that basic residues at P1 and Arg at P2 interact with the acidic residues E63 and D24 in both molecules, and with Tyr at Pc sitting in a hydrophobic pocket with the hydroxyl interacting with D116 of B15, while the hydrophobic amino acids at Pc interacting with a more hydrophobic pocket in B19.

More recently, isolation of class I molecules from a B19 cell line followed by HPLC and mass spectrometric analysis of single peptides (LC-MS/MS, or immunopeptidomics) was performed (18), which confirmed and extended the previous results. As this cell line expressed BF1 molecules at a higher level than is found on normal cells, an analysis of the peptides with Arg at P2 (almost certainly from the BF2 molecule) was performed (Fig. 1E). Of the 896 peptides, amino acids at P1 were over 25% Lys, 17% Val and 15% Ile, with lesser amounts of Arg, Gln and Thr and then Ala, Leu, Met and Ser. Over 50% of all peptides had either Phe or Tyr at P3, with around 15% Leu and lesser amounts of other amino acids. At P5, 19% Ser, 14% Pro, 11% Gly, 9% Ala and other amino acids at lower amounts were found. Around 82% of amino acids at Pc were Phe, Ile, Leu, Pro, Val or Tyr, although which predominated depended on the length of peptide, for which there were 328 8mers, 282 9mers, 109 10mers and 51 11mers (totaling 770 of the 896 peptides, with nearly all of the rest being longer).

### The structural overview of BF2*1901 is similar to BF2*1501

On the basis of the motifs determined above, peptides from MDV that might bind different chicken MHC molecules were predicted (S1 Table, S2 Table) (22). The structure of chicken MHC I molecule BF2*1901 complexed with predicted MDV peptide RY8 was determined to resolution of 2.0 Å with two molecules in one asymmetric unit, while the structure of BF2*1901 with influenza H5N1 virus M1 peptide IL9 was determined at 2.0 Å with one molecule in an asymmetric unit. The overall structure of BF2*1901 retains the common characteristics of MHC class I molecules from other vertebrates including chickens: the extracellular region of the BF2*1901 heavy chain folds into three different domains (Fig. 2A); the α1 and α2 domains form a typical MHC I peptide binding groove (PBG), which contains two α-helices and eight β-strands; the RY8 or IL9 peptide lies along the PBG, as shown through well-determined electron density maps (Fig. 2B,C). The α3 domain of BF2*1901 and β_2_m are typical immunoglobulin superfamily domains and underpin the α1 and α2 domains. The superposition of BF2*1901/RY8 onto the structure of BF2*1901/IL9 generated a root mean square deviation (RMSD) of 0.533 Å. This superposition of the two structures showed that the most distinct portion of the two molecules to be located in the middle of the α2 helix, which covers residues 145E to 149Y (Fig. 2B). For BF2*1901/IL9, the loop at the middle of α2 helix is closer to the C terminal of the peptide helix, compared to the structure of BF2*1901/RY8.

**Fig 2.**
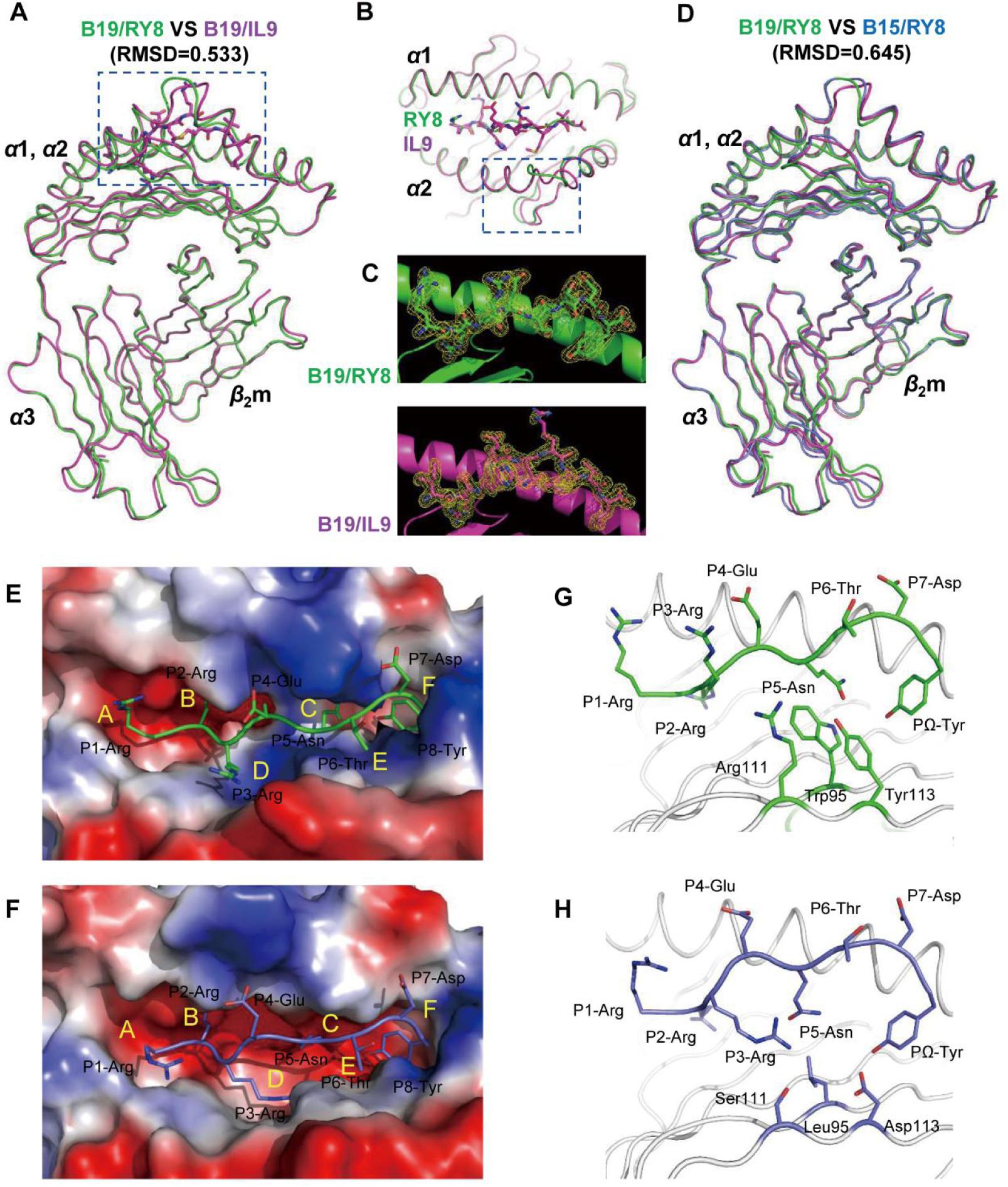
Structural overview of BF2*1901 and its shallow and narrow peptide binding groove compared to BF2*1501. A, Superimposed overall structures of B19 complexed to MDV peptide RY8 (green), and influenza H5N1 virus M1 peptide IL9 (purple). The RMSDs of the monomer MHCs were determined to be 0.533 Å for B19/RY8 VS B19/IL9. B, The alignment of α1α2 domain of B19/RY8 and B19/IL9 indicates a conformational difference at the α2 helices of the two structures. C, The peptide RY8 (green) and IL9 (purple) in the structures of B19 are presented with the 2Fo-Fc electron density maps at the 1.0 σ contour level. D, Superimposed overall structures of B19/RY8 (green) and B15 complexed to the similar MDV peptide RY8 (PDB: 6LHH, blue). The RMSDs of the monomer MHCs were determined to be 0.645 Å for B19/RY8 VS B15/RY8. E, The vacuum electrostatic surface potential shows the peptide binding groove of B19 complexed to MDV peptide RY8 (green). F, The vacuum electrostatic surface potential shows the peptide binding groove of B15 complexed to MDV peptide RY8 (PDB: 6LHH, Blue). G, The peptide RY8 (green) in B19 is shown in green cartoon and three large B19-specific residues at the bottom of the groove, i.e. Trp95, Arg111, Tyr113 are show in green sticks. H, The peptide RY8 in B15 (PDB: 6LHH) is shown in blue cartoon and three B15-specific residues at the bottom of the groove, i.e. Leu95, Ser111, Asp113 with short side chains are show in blue sticks.

The overall structures of BF2*1901 are extremely similar to those of BF2*1501 (Fig. 2D). The superposition of BF2*1901/RY8 onto the previously determined structure of BF2*1501 complexed to the same peptide RY8 generated an RMSD of 0.645 Å. Moreover, the identity of the amino acid sequences of BF2*1901 and BF2*1501 is 97.04% (S4 Fig.). As expected, only two polymorphic residues S69T and I79T (for BF2*1901 versus BF2*1501) are located in the α1 and α2 domains. The structural comparison highlights the altered solvent exposure of residue I79T, which may have important role in the distinct MHC restrictions for T-cell receptor (TCR) recognition. For the conformation of the main chain of BF2*1901 and BF2*1501, BF2*1901/RY8 has a similar conformation as BF2*1501 at the middle of the α2 helix, while BF2*1901/IL9 has a conformational shift at this place when compared to BF2*1501 (Fig. 2D).

### The shallow and narrow peptide binding groove of BF2*1901

Like most mammalian classical class I molecules including BF2*1501, BF2*1901 has obvious pockets A–F (Fig. 5E). However, only the pockets A and B of BF2*1901 are very similar to BF2*1501, while the C, D, E and F pockets in the PBG of BF2*1901 possess their own allele-specific features. Pockets A of BF2*1501 and BF2*1901 present an open and large space with a relative negative charge to accommodate P1 residue at the N terminus of peptide. Furthermore, the Pocket B for both BF2*1501 and BF2*1901 are very deep and negatively-charged. The conserved salt bridge between the P2-Arg of the peptides and residues Asp24, Thr34, and Glu62 of the main chains of both BF2*1501 and BF2*1901 can be observed (S5 Fig.).

In contrast, the major distinct portions of the PBG of BF2*1501 and BF2*1901 locate to the C, D, and E pockets in the center of the groove. Compared to BF2*1501, BF2*1901 has a much narrower and shallower groove (Fig. 2E,G). The main-chain atoms of the α1/α2 platform of BF2*1901 and BF2*1501 are nearly superimposable (Fig. 2D), so that the narrow groove of BF2*1901 and the wide groove of BF2*1501 are due entirely to the different side chains of amino acids pointing into the groove. In particular, the large residues W95, R111 and Y113 from the β-strands on the bottom of PBG of BF2*1901 are replaced by the much smaller L95, S111 and D113 in BF2*1501 (Fig. 2F,H). These large overhanging residues with bulky side chains occupies most space of the C, D, and E pockets in PBG of BF2*1901. The distances of the bottom of the PBG to the binding peptide (represented by the upper atom of the side chain of W95, R111 and Y113 to the corresponding Cα-atom of P4, P5, and P6 residues of the peptides) are 4.76 Å, 4.89 Å and 6.59 Å, compared to the longer distances in BF2*1501, i.e. 7.00 Å, 10.16 Å, and 9.16 Å. Thus, the deep and wide middle portion of PBG of BF2*1501 allows the remodeling of the groove to accept peptides with promiscuous secondary anchor residues in the middle and adopt various conformations.

### The tight but flexible P1 anchor of BF2*1501 compared to BF2*1901

In the structure of BF2*1501 and BF2*1901, the conserved residue Glu65 enables the A pocket to be relatively negatively-charged, as it is in HLA-B27 (45). Thus, the peptides with positive charged P1 residues are preferred by both BF2*1501 and BF2*1901(15, 21). However, the detailed superposition of the two chicken MHC I showed different modes of P1 anchoring. We superposed A pockets of the BF2*1901/RY8 and all the available structures of BF2*1501 complexed to peptides with positive P1 anchors (Fig. 3A). The superposition clearly shows the similar conformation of P1-Arg in the two molecules M1 and M2 of the asymmetrical unit of B19/RY8 structure (Fig. 3A), but the P1-Arg of flu peptide PA124 presented by BF2*1501 is closer to the α1 helix (Fig. 3A) while the P1-Arg of peptides RY8 and chicken calcium-binding protein peptide CBP in BF2*1501 structure is closer to the peptide itself (S6A Fig.). Moreover, in the M1 of BF2*1901/RY8, the hydrogen bond between P1-Arg of peptide RY8 and the Glu65 of BF2*1901 is 3.58 Å (Fig. 3B), while no interaction between them is observed in B19/RY8 M2 (not shown). In contrast, the closer and higher binding with two hydrogen bonds of between P1-Arg in B15/PA124 and the residues Tyr61 (2.89 Å) and Glu65 (2.90 Å) in α1 helix of B15 can be observed (Fig. 3C).

**Fig 3.**
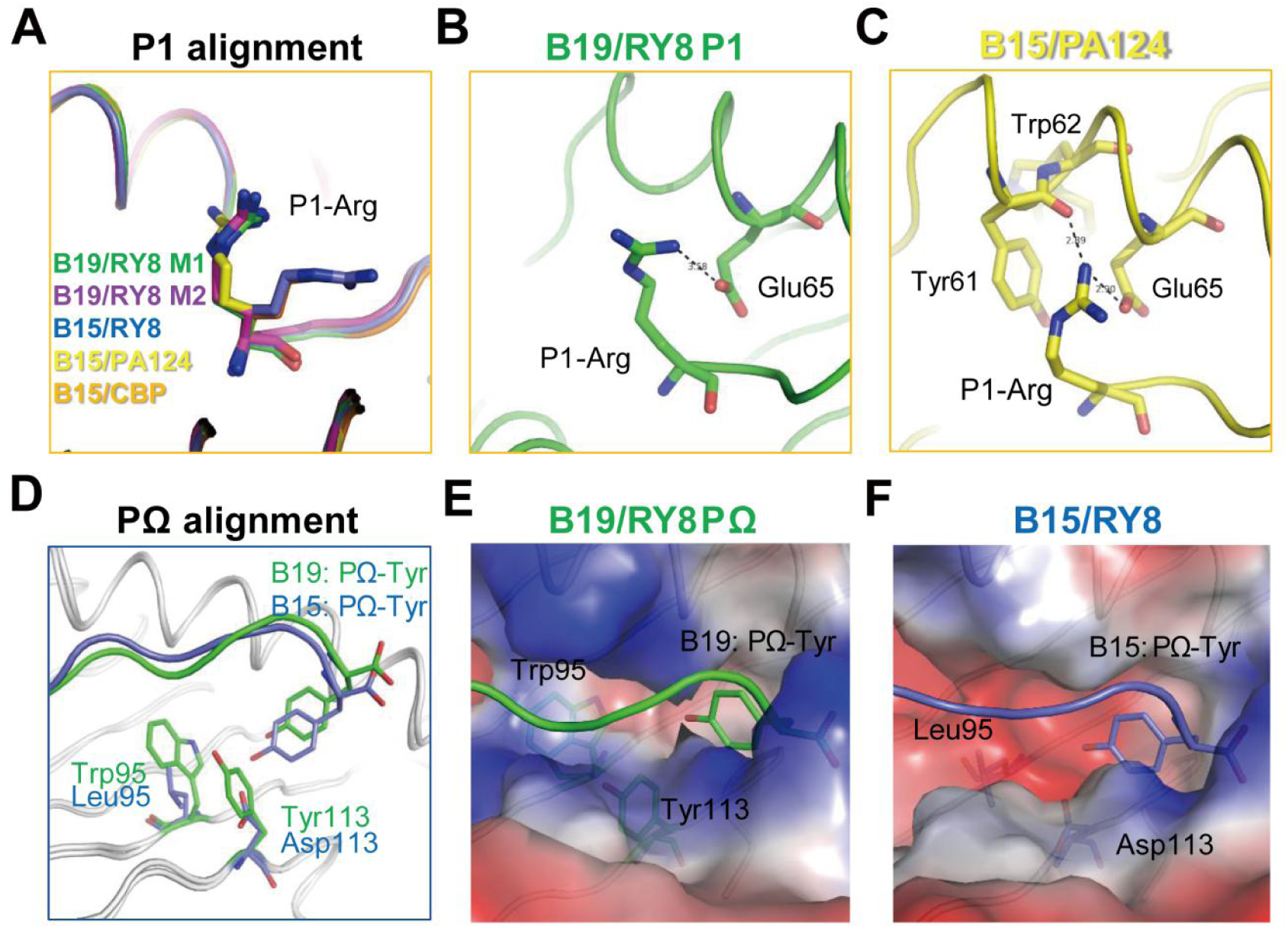
The different features of A and F pockets of BF2*1901 compared to BF2*1501. A, The superposing between A pockets of the two molecules M1 (green) and M2 (purple) of the asymmetrical unit of B19/RY8 structure, and also the B15 complexed to MDV peptide RY8 (PDB: 6LHH, blue), flu peptide PA124 (PDB: 6IRL, yellow) and chicken calcium-binding protein peptide CBP (PDB: 6KX9, orange). The superposition clearly shows the similar conformation of P1-Arg in B19 M1 and M2. But the P1-Arg of B15/PA124 is closer to the α1 helix. While the P1-Arg of peptides RY8 and CBP are closer to the peptide itself. B, The weak hydrogen bond between P1-Arg of peptide RY8 in B19/RY8 M1 (green) and the Glu65 of B19. No interaction between them is observed in B19/RY8 M2 (not shown). C, The closer binding between P1-Arg in B15/PA124 (yellow) and the residues Tyr61 and Glu65 in α1 helix of B15. D, The superposing between B19/RY8 and B15/RY8 according to the α-C of α1α2 domains. The superposition clearly shows the higher position of PΩ-Tyr of peptide RY8 in B19/RY8 structure, compared to the PΩ-Tyr of peptide RY8 in B15/RY8 structure. E and F, The vacuum electrostatic surface potential shows the narrow and shallow F pocket of B19 (E, peptide RY8 in green sticks) compared to the B15/RY8 (PDB: 6LHH) (F, peptide RY8 in blue sticks).

### The narrow and shallow F pocket of BF2*1901

Then we focused on the PΩ anchors, for which BF2*1501-binding peptides strongly prefer Tyr, but BF2*1901-binding peptides have a variety of hydrophobic anchor resides. When we superposed the structures of B19/RY8 and B15/RY8 according to the Cα of α1/α2 domains, we found that the position of PΩ-Tyr of peptide RY8 in B19/RY8 structure is higher compared to RY8 in B15/RY8 structure (Fig. 3D). The detailed analysis of the F pockets of BF2*1901 shows the narrow and shallow F pocket which is occupied by the residues Trp95 and Tyr113 with large side chains (Fig. 3E). Thus, the two B19-specific residues Trp95 and Tyr113 work like two bricks to bolster up the PΩ-Tyr of peptide RY8 in B19. In contrast, residues Leu95 and Asp113 in the F pocket of B15/RY8 make room for the deep location of PΩ-Tyr of peptide RY8 in B15 (Fig. 3F). Furthermore, the relatively smaller F pocket of BF2*1901 can accommodate the peptide IL9 with PΩ-Leu (S6B Fig.).

### The flexible but tight binding of P3 anchor of BF2*1501 compared to BF2*1901

When we compared the conformation of the same peptide RY8 presented by BF2*1901 and BF2*1501, we found the P3-Arg had distinct conformation within the two structures. The P3-Arg protrudes the side chain out of the D pocket of B19/RY8 groove (Fig. 4A), while in B15/RY8 structure, the P3-Arg anchor puts the side chain into the D pocket of B15/RY8 groove (Fig. 4B). Furthermore, we aligned available BF2*1901 and BF2*1501 structures, and found the P3 anchor of peptides presented by BF2*1501 can accommodate different conformations with the side chains pointing into or outside the D pocket. In contrast, the P3 anchors of peptides presented by BF2*1901 all protrude out of the D pocket (Fig. 4C). Further analysis found the shallow and narrow D pocket of B19/RY8 groove is occupied by the large positive charged residue Arg111 (Fig.4D,F), in contrast to the large D pocket of BF2*1501 with the small residue Ser111 (Fig. 4E,G). In the structures of B19/RY8 and B19/IL9, P3-Arg (Fig. 4D) and P3-His (Fig. 4F) protrude their side chains out of the D pockets. Meanwhile, a π-π interaction can be observed between P3-His of peptide IL9 and the residue Tyr156 of BF2*1901 (Fig. 4F), which may partly compensate the week binding of P3 anchor out of the BF2*1901 groove. In the structure of B15/RY8, the P3-Arg locates in the larger D pocket and the hydrogen bond between P3-Arg and Ser111 of B15 can be observed. In contrast, the P3-Glu points out of the D pocket in the groove of BF2*1501/PA124 due to the presence in the D pocket of P5-His of peptide PA124. These analyses indicate a flexible but tight binding of P3 anchor of BF2*1501 compared to BF2*1901.

**Fig 4.**
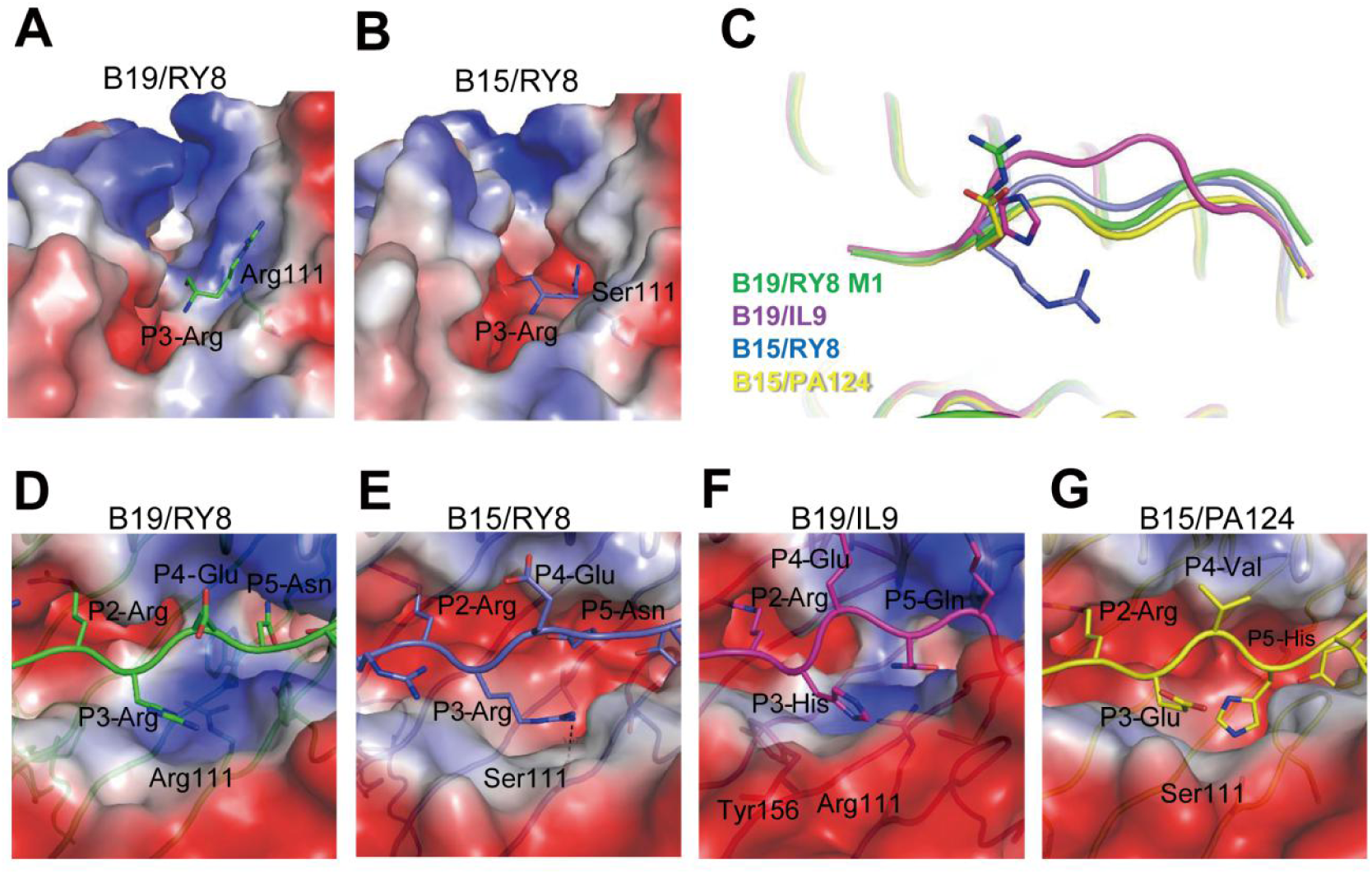
The flexible but tight binding of P3 anchor of BF2*1501 compared to BF2*1901. A, The P3-Arg anchor protrudes its side chain out of the shallow and narrow D pocket of B19/RY8 groove, which is occupied by the large positive charged residue Arg111. B, The P3-Arg anchor puts its side chain into the D pocket of B15/RY8 (PDB: 6LHH) groove, where small residue Ser111 locates. C, The superposing of B19/RY8 (green), B19/IL9 (purple), B15/RY8 (PDB: 6LHH, blue) and B15/PA124 (PDB: 6IRL, yellow). The superposition clearly shows two different conformations of P3 anchors of peptides presented by B15, i.e P3-Arg in RY8 and P3-Glu in PA124. D, The smaller D pocket of B19/RY8. E, The larger D pocket of B15/RY8 and the hydrogen bond between P3-Arg and Ser111 of B15. F, The similar D pocket and P3 conformation of B19/IL9 as in B19/RY8. G, The P5-His of peptide PA124 occupies the D pocket of B15/PA124. Thus, P3-Glu points out of the D pocket.

### The higher middle portion of α2 helix of BF2*1901/RY8 for TCR docking

In addition to the detailed analysis of the peptide anchoring of BF2*1901 and BF2*1501, we also analyzed the MHC heavy chain itself in these two closely-related chicken MHC I molecules. The superimposition of B19/RY8 and B15/RY8, according to the Cα of α1/α2 domains showed different conformations of the middle potion in the α2 helices of the two structures (Fig. 5A). The middle portion of α2 helix of B19, covering Trp144 to Tyr 149 had a higher position compared to the corresponding residues of B15. The structure analysis showed that the two larger residues Arg111 and Tyr113 from the β-sheet of BF2*1901 jack up the α2 helix through the interaction with Tyr149 and Trp144 of α2 helix (Fig. 5B). In contrast, the α2 helix of B15 touches down due to the short sidechains of residues Ser111 and Asp113 in the β sheet (Fig. 5C). Interestingly, in the two structures of B19 we determined here, the middle potion in the α2 helix of B19/IL9 shows a distinct conformation compared to the corresponding position of B19/RY8 (Fig. 5D). The superposing of the two structures showed that the different secondary anchor residue of peptides RY8 and IL9 lead to the conformation shift of the α2 helices in the two structures. The large residue P7-Met of peptide IL9 pushes the Tyr149 out of the peptide binding groove, which is difference for the residue P6-Thr of peptide RY8 (Fig. 5E-G). The middle portion in the α2 helix of MHC I locates at the highest position or so-called super-bulged conformation in the TCR docking surface(46, 47). Thus, the conformational specificity of BF2*1901 at this region may lead to uncommon TCR docking strategy, which may imply a limited TCR repertoire of BF2*1901.

**Fig 5.**
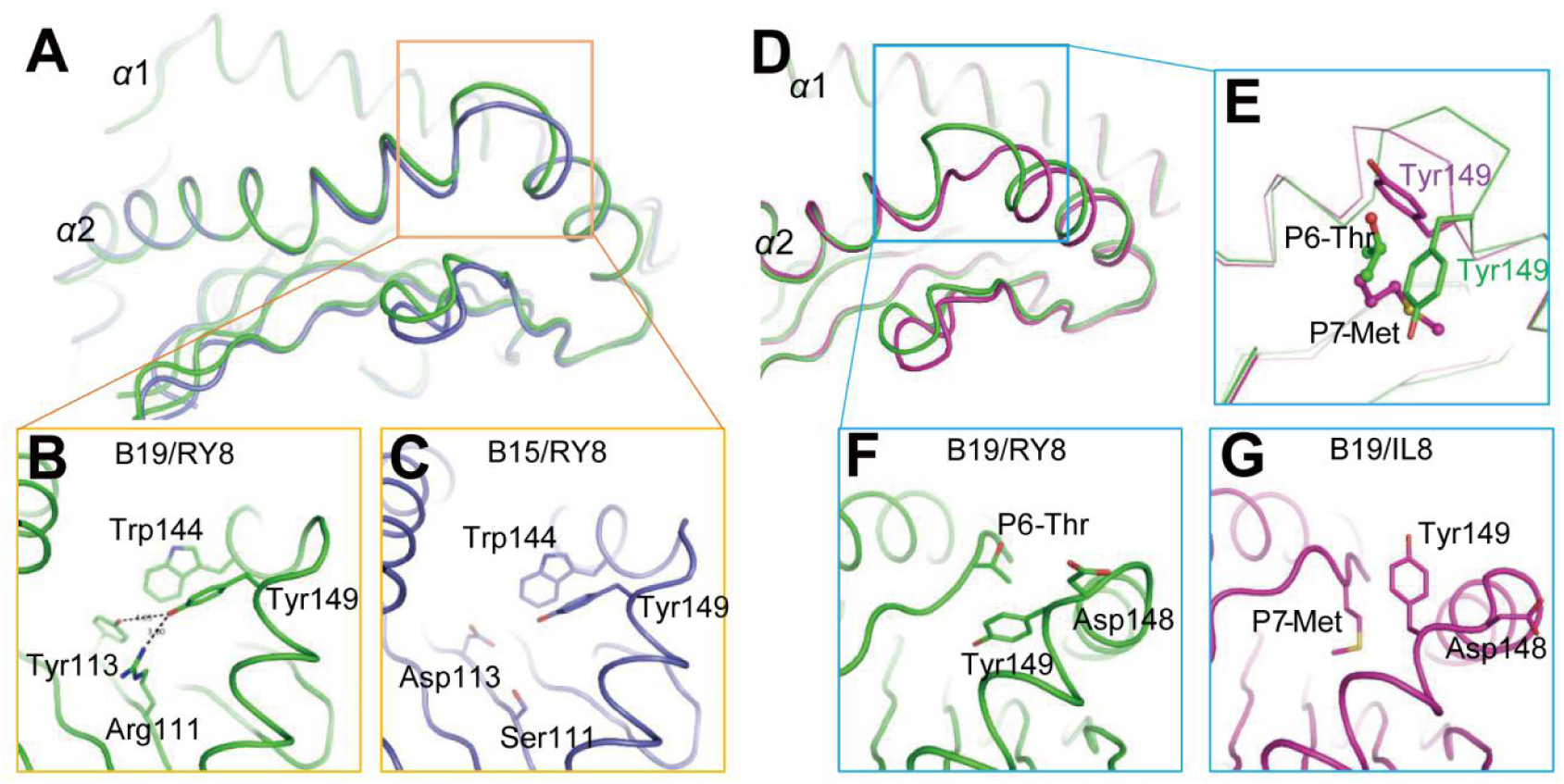
The higher α2 helix of BF2*1901/RY8 compared to BF2*1501/RY8 and BF2*1901/IL9. A, Superimposed structures of B19/RY8 (green) and B15/RY8 (PDB: 6LHH, blue), according to the α-C of α1α2 domains. The different conformations of the middle potion in the α2 helices were shown in yellow square. B, The two larger residues Arg111 and Tyr113 jack up the α2 helix of B19 through the interaction with Tyr149 and Trp144. C, In B15/RY8 (PDB: 6LHH, blue), the α2 helix touches down due to the short sidechains of residues Ser111 and Asp113. D, Superimposed B19/RY8 (green) and B19/IL9 (purple) according to the α-C of α1α2 domains. The different conformations of the middle potion in the α2 helices were shown in blue square. E, In the structural comparison between B19/RY8 (green) and B19/IL9 (purple), the large residue P7-Met of peptide IL9 pushes the Tyr149 out of the peptide binding groove, which is difference for the residue P6-Thr of peptide RY8. F and G, The detailed conformational comparison between of B19/RY8 and B19/IL9.

## DISCUSSION

The correlation of resistance to Marek’s disease with peptide repertoires for chicken class I (BF2) molecules is very clear(15, 18, 19, 22, 26), but the reasons why the B19 haplotype confers more susceptibility than the B15 haplotype and the cell surface class I level of B19 cells is higher than B15 cells, and whether the BF2*1901 molecule has more fastidious peptide-binding than BF2*1501 have all remained unclear. In this paper, we confirm that the viral loads after MDV infection of B19 chickens are much higher than of B15 chickens, describe and compare the detailed peptide motifs from B19 cells versus B15 cells, and show by multiple crystal structures how the narrow and shallow peptide-binding groove of BF2*1901 molecules results in a less promiscuous binding than the relative larger and deeper groove of BF2*1501.

We have recently derived chicken lines with various MHC haplotypes and examined some of them (including the line bearing the B19 haplotype) for response to MDV (43). Here we use RT-qPCR to show that the basal levels of various cytokines are generally similar in the lines with B15 and B19, but with higher active cytokine IFN-γ and lower inhibitory cytokine IL-10 in B15 chickens. Furthermore, the virus levels determined by qPCR after MDV infection begin to differ sharply at 12 dpi. Much more virus is found in B19 chickens, in agreement with the published hierarchy of susceptibility to Marek’s disease (12).

We also present detailed evidence for the self-peptides bound to class I molecules presented by B19 cells: sequences from individual peptides and peptide pools from blood and spleen cells by gas phase sequencing, as well as peptides with Arg at P2 from an MDV-transformed cell line by immunopeptidomics. These results confirm and extend the B19 class I motif originally described(21), but they fail to explain in any obvious way the relative MDV susceptibility of B19 compared to B15 chickens in terms of peptide repertoire. In comparison with the sequences of individual peptides and peptide pools from peptides of B15 cells presented previously(15, 21), both molecules require an Arg at P2, but B15 prefers a basic residue at P1 and a Tyr at Pc compared to multiple amino acids found for B19 at both positions. Thus, the dominantly-expressed class I molecule of B19 seems more promiscuous based on the peptide sequences than the class I molecule of B15, which is the opposite of what has been seen up to now in terms of MDV resistance(18, 19).

We resolve this conundrum using structures of B15 and B19 class I molecules (BF2*1501 and BF2*1901) bound to multiple peptides, including the same peptide (RY8) bound to both molecules. Although both B15 and B19 molecules bind the amino terminus of the peptide in pocket A and require Arg as the anchor residue at P2 in a deep pocket B containing Asp24, B19 molecule has many larger residues leading to an overall narrower and shallower peptide-binding groove in pockets C, D, E and F. The larger Trp95 and Tyr113 of BF2*1901 leads to a much narrower and shallower pocket F than Leu95 and Asp113 of BF2*1501. Thus, the various amino acids found at the C-terminal anchor residue of BF2*1901 are likely bound with much less affinity (with therefore likely fewer total peptides) than the Tyr overwhelmingly favored by BF2*1501. Similarly, the larger Arg111 residue of BF2*1901 leads to a much narrower and shallower pocket D than Ser111 of BF2*1501. The side chains of residues at P3 are all forced out of the groove in BF2*1901, whereas most are accommodated (as so-called secondary anchors) in larger pocket D of BF2*1501, with one exception due to the peptide residue at P5 occupying pocket D. Thus, the location of the middle of the peptide is higher out of the groove, with again likely less affinity and fewer numbers of peptides for BF2*1901. Also, Trp95 in BF2*1901 is much larger than Leu95 in BF2*1501, so that pocket C is similarly affected, as shown by the deeper anchoring of P5-Asn of peptide RY8 in BF2*1501 compared to P5-Asn of peptide RY8 in BF2*1901 (Fig. 2G,H). Thus, the wider and deeper peptide-binding groove of BF2*1501 means that more peptides can be bound, as opposed to BF2*1901 for which only fewer peptides with the highest affinity will bind. Meanwhile, our BF2*1901 structure showed a π-π interaction between P3-His of peptide IL9 and the residue Tyr156 of BF2*1901 (Fig. 6F), which partly compensates the week binding of P3 anchor out of the BF2*1901 groove. This may explain the why >50% BF2*1901-binding peptides prefer Phe or Tyr as P3 anchor based on the immunopeptidomic data.

Despite both molecules having Glu65 which could bind to basic residues at peptide position P1 (as in the human class I molecule HLA-B27,(45), the conformation of the amino acids at P1 of peptides varied considerably, even for the peptides bound to BF2*1501 which all have basic amino acids at P1. The basic sidechain of P1-Arg from only one peptide PA124 bound to BF2*1501 formed a salt bridge (Fig. 3C) with Glu65 (2.90 Å) and a hydrogen bond with Tyr61 (2.89 Å). While although in molecule 1 of BF2*1901/RY8 asymmetric unit, sidechain of P1-Arg from peptide RY8 bounds to Glu65 of BF2*1901 with a weak hydrogen bond (3.58 Å), no interaction of P1-Arg with residue of BF2*1901 can be observed in molecule 2 of BF2*1901/RY8. The reason for these differences remains mysterious, but it still indicates a tight anchoring of P1 residue of BF2*1501-binding peptide compared to the ones of BF2*1901.

Finally, the larger residues Arg111 and Tyr113 from the β-sheet interact with Tyr149 and Trp144 of the α2 helix to raise the middle of the α2 helix in BF2*1901, compared to BF2*1501 which has Ser111 and Asp113. Moreover, conformational changes based on the sequence of the peptide bound to BF2*1901 can raise the α2 helix even further, with Met but not Thr at P7 pushing Tyr149 out of the groove, leading to a super-bulged surface which most T cell receptors may not bind (46). Although previously it has been speculated that the peptide repertoire of MHC molecules may affect the T cell repertoire(18, 19), this observation of a fastidious class I molecule that may not be easily recognized by most T cell receptors provides a new mechanism by which this situation might occur.

In summary, we provide further evidence that B19 chickens are more susceptible to MDV than B15 chickens, we conduct the first detailed analysis of self-peptides leading to the peptide motif of BF2*1901, and we present several structures of BF2*1501 and BF2*1901. We find that the self-peptides bound to BF2*1901 may appear more various than those of BF2*1501, but that the structures show the wider and deeper peptide-binding groove of BF2*1501 means that it will accept many more peptides overall. This finding is consistent with the width and depth of the whole range of promiscuous to fastidious molecules(18, 22, 25, 26), and suggests that the peptides found bound to BF2*1901 are the very best binders to a narrow and shallow groove, in which many parts of the peptide binding are important, not just the particular amino acids in the positions of the anchor residues. Moreover, the higher position of the α2 helix of BF2*1901, particularly when affected by particular peptide residues, compared to BF2*1501 suggests another level, that of T cell repertoire, which generalist and specialist alleles may affect.

## MATERIALS AND METHODS

### Animals

Two chicken populations with B15 and B19 haplotypes were bred and maintained at National Poultry Laboratory Animal Resource Bank, which is affiliated with Harbin Veterinary Research Institute, Chinese Academy of Agricultural Sciences. The specific pathogen-free (SPF) chickens were kept in positive pressure isolators with high efficiency particulate air (HEPA) filters throughout life and have been free from the 19 avian diseases which conform to the request stipulated by national standard of GB 17999.1-2008 *SPF chicken-Microbiological surveillance-Part 1: General rules for the microbiological surveillance for SPF chicken* for 15 generations. The environment index of the breed facility conforms to the standard of GB 14925-2010, with ^60^Co-sterilized feed and acidified drinking water; the laboratory animal production license issued by local government is SCXK (HEI) 2006-009. Both the original breeding populations were selectively bred and maintained for 5 generations based on the genomic sequences of *BF1, BF2, BLB1, BLB2* and *BG4* genes(43). The research was approved by Committee on the Ethics of Animal Experiments of Harbin Veterinary Research Institute (HVRI) of the Chinese Academy of Agricultural Sciences (CAAS).

### Expression levels of IFN-γ, IL-18, and IL-10 mRNA of B15 and B19 chickens

Four duplex TaqMan probe real-time fluorescence quantitative RT-PCR (dqRT-PCR) methods for chicken IFN-γ, IL-18, and IL-10, respectively, were established as previously described(48–50).

### The chicken infection by MDV Md 5 and the dynamic viral loads

Fresh or rejuvenated MDV Md5-infected chicken peripheral blood was diluted by 50 times with DMEM, and 1-day-old B19 chickens (n=9) and B15 (n=9) chickens were inoculated intraperitoneally (i.p.) with 500 µL each. Control animals (n=5 for each line) were inoculated with 500 µL DMEM. The feather pulps of 3 chickens in each group were collected randomly at 4, 7, 9, and 12 dpi. Virus copy number was detected by dqPCR using the MDV *meq* gene to detect the virus load, and the chicken egg iron transfer protein gene (ovo) as an internal reference.

### Sequencing of peptides bound to class I molecules

As described in detail (15, 18), monoclonal antibodies to chicken class I heavy chain (F21-2) and to chicken β_2_-microglobulin (F21-21) were used to isolate class I molecules from detergent lysates of red blood cells, spleen cells or the B19 cell line MDCC-265L. The peptides from the *ex vivo* cells were separated by reverse-phase HPLC using a Pharmacia SMART system, with sequencing of individual peptide peaks or of whole peptide pools using an Applied Biosystems 475A gas phase sequencing. The peptides from the cell line were analyzed by LC-MS/MS using the Q-Exactive (Thermo Scientific) and TripleTOF 5600 (AB Sciex) systems.

### Preparation of BF2*1901 protein

The cDNAs for BF2*1901 (GenBank: Z54317.1) were synthesized (Genewiz Inc, Beijing). The extracellular domain (residues 1–270) of BF2*1901 were cloned into a pET21a vector (Novagen) and transformed into *Escherichia coli* strain BL21 (DE3). Dilution-renaturation and purification of BF2*1901 molecules assembled with chicken β_2_m (chβ_2_m) (expressing residues 1-98) (26) and peptides were performed as described previously(51).

### X-ray Crystallography, Structure Determination, and Refinement

Crystallization of was performed using the sitting drop vapor diffusion technique. BF2*1901/RY8 crystals were observed in 0.15 M KBr and 30% w/v polyethylene glycol monomethyl ether 2000 at a protein concentration of 13.5 mg/mL. Single crystals of BF2*1901/IL9 were grown in 0.1 M BIS-TRIS pH 6.5 and 28% w/v polyethylene glycol monomethyl ether 2000 at a protein concentration of 12.5 mg/mL. Diffraction data for both crystals were collected at 100 K at the SSRF BEAMLINE BL17U, Shanghai, China. The collected intensities were subsequently processed and scaled using the DENZO program and the HKL2000 software package (HKL Research)(52). The structure of BF2*1901 was determined by molecular replacement using BF2*1201 (Protein Data Bank [PDB] code 5YMW) as a search model in the Crystallography & NMR System (CNS) (53) and COOT (54), refined with REFMAC5 (55) and PHENIX (56) and assessed with PROCHECK (57) (S1 Table). Structure-related figures were generated using PyMOL (http://www.pymol.org/). The sequence alignment was generated with Clustal X (58) and ESPript(59).

### Accession numbers

Protein Data Bank (http://www.rcsb.org) accession codes are 7WBG for BF2*1901/RY8 and 7WBI for BF2*1901/IL9.

## ACNOWLEDGMENTS

This work was supported by the National Natural Science Foundation of China (NSFC) (81971501). W.J.L. is supported by the Excellent Young Scientist Program of NSFC (81822040). D.W.A., H.-J.W. and J.K. (and Jan Salomonsen, sadly deceased and much missed) were supported by F. Hofmann-La Roche while working at the Basel Institute for Immunology. The work by V.N., W.M. and N.T. was supported by the BBSRC at the Pirbright Institute at Compton. J.K. is supported by an Investigator Award from the Wellcome Trust (110106/Z/15/Z).

## SUPPLEMENTARY INFORMATION

**S1 Fig. Dual qPCR amplification and standard curves for target gene and reference gene plasmids.**

Amplification curve and standard curve for IFN-γ/28s, IL-18/28s rRNA, and IL-10/28s rRNA, respectively; FAM: target gene plasmid amplification; HEX: reference gene amplification.

**S2 Fig. The relative background cytokine expression in B15 and B19 SPF chickens.**

The relative expression levels of IL-10 (A), IFN-r (B), and IL-18 (C) inperipheral blood were detected with dqRT-PCR method. The data came from 11 healthy B15 chickens and 6 of B19 chickens at different days old.

**S3 Fig. Comparison of the relative expression levels of cytokines in different tissues of B15 and B19 at 70d-old.**

The relative expression levels of IFN-γ, IL-18, and IL-10 in lung with respiratory tract, thymus, bursa of Fabricius, spleen and peripheral blood were detected with dqRT-PCR method. The data came from each 3 of 70 days-old B15 and B19 chickens.

**S4 Fig. Structure-based sequence alignment of BF2*1901 and BF2*1501.**

Cylinders indicate α-helices, and black arrows indicate β-strands. Residues highlighted in red are completely conserved, and residues in blue boxes are highly (>80%) conserved. Residues that play a critical role in the conformations of Mamu-A*02-presented peptides are marked with deep blue asterisks. The sequence alignment was generated with Clustal X (58) and ESPript(59).

**S5 Fig. Comparison of the B pocket of the BF2*1501 and BF2*1901 to accommodate peptide RY8.**

A, Structure of B pocket in B19/RY8 (green). B, B pocket of B15/RY8 (PDB: 6LHH, blue). The hydrogen bond between P2-Arg of peptide RY8 in B19/RY8 and B15/RY8 are shown in black dashed lines.

**S6 Fig. The A pocket of B15/RY8 and F pocket of BF2*1901/IL9.**

A, The intra chain hydrogen bond of P1-Arg in the B15/RY8 (Blue). B, The PΩ-Leu of peptide IL9 in BF2*1901/IL9 structure inserts its side chain into Pocket F of BF2*1901. The vacuum electrostatic surface potential shows the narrow and shallow F pocket of B19 with peptide IL9 in purple sticks.

## REFERENCES

1. Goodnow CC. 2021. COVID-19, varying genetic resistance to viral disease and immune tolerance checkpoints. Immunol Cell Biol 99:177–191.

2. Kenney SP, Wang Q, Vlasova A, Jung K, Saif L. 2021. Naturally occurring animal coronaviruses as models for studying highly pathogenic human coronaviral disease. Vet Pathol 58:438–452.

3. Reagan RL, Brueckner AL. 1952. Electron microscope studies of four strains of infectious bronchitis virus. Am J Vet Res 13:417–8.

4. Bertzbach LD, Conradie AM, You Y, Kaufer BB. 2020. Latest Insights into marek’s disease virus pathogenesis and tumorigenesis. Cancers (Basel) 12.

5. Osterrieder N, Kamil JP, Schumacher D, Tischer BK, Trapp S. 2006. Marek’s disease virus: from miasma to model. Nat Rev Microbiol 4:283–94.

6. Reddy SM, Izumiya Y, Lupiani B. 2017. Marek’s disease vaccines: Current status, and strategies for improvement and development of vector vaccines. Vet Microbiol 206:113–120.

7. Smith J, Lipkin E, Soller M, Fulton JE, Burt DW. 2020. Mapping QTL associated with resistance to avian oncogenic marek’s disease virus (MDV) reveals major candidate genes and variants. Genes (Basel) 11.

8. Vallejo RL, Bacon LD, Liu HC, Witter RL, Groenen MA, Hillel J, Cheng HH. 1998. Genetic mapping of quantitative trait loci affecting susceptibility to Marek’s disease virus induced tumors in F2 intercross chickens. Genetics 148:349–60.

9. Wolc A, Arango J, Jankowski T, Settar P, Fulton JE, O’Sullivan NP, Fernando R, Garrick DJ, Dekkers JC. 2013. Genome-wide association study for Marek’s disease mortality in layer chickens. Avian Dis 57:395–400.

10. Briles WE, Briles RW, Taffs RE, Stone HA. 1983. Resistance to a malignant lymphoma in chickens is mapped to subregion of major histocompatibility (B) complex. Science 219:977–9.

11. Briles WE, Stone HA, Cole RK. 1977. Marek’s disease: effects of B histocompatibility alloalleles in resistant and susceptible chicken lines. Science 195:193–5.

12. Plachy J, Pink JR, Hála K. 1992. Biology of the chicken MHC (B complex). Crit Rev Immunol 12:47–79.

13. Miller MM, Taylor RL, Jr. 2016. Brief review of the chicken major histocompatibility complex: the genes, their distribution on chromosome 16, and their contributions to disease resistance. Poult Sci 95:375–92.

14. Kim T, Hunt HD, Parcells MS, van Santen V, Ewald SJ. 2018. Two class I genes of the chicken MHC have different functions: BF1 is recognized by NK cells while BF2 is recognized by CTLs. Immunogenetics 70:599–611.

15. Wallny HJ, Avila D, Hunt LG, Powell TJ, Riegert P, Salomonsen J, Skjødt K, Vainio O, Vilbois F, Wiles MV, Kaufman J. 2006. Peptide motifs of the single dominantly expressed class I molecule explain the striking MHC-determined response to Rous sarcoma virus in chickens. Proc Natl Acad Sci U S A 103:1434–9.

16. Goto RM, Wang Y, Taylor RL, Jr., Wakenell PS, Hosomichi K, Shiina T, Blackmore CS, Briles WE, Miller MM. 2009. BG1 has a major role in MHC-linked resistance to malignant lymphoma in the chicken. Proc Natl Acad Sci U S A 106:16740–5.

17. Rogers SL, Kaufman J. 2008. High allelic polymorphism, moderate sequence diversity and diversifying selection for B-NK but not B-lec, the pair of lectin-like receptor genes in the chicken MHC. Immunogenetics 60:461–75.

18. Chappell P, Meziane el K, Harrison M, Magiera Ł, Hermann C, Mears L, Wrobel AG, Durant C, Nielsen LL, Buus S, Ternette N, Mwangi W, Butter C, Nair V, Ahyee T, Duggleby R, Madrigal A, Roversi P, Lea SM, Kaufman J. 2015. Expression levels of MHC class I molecules are inversely correlated with promiscuity of peptide binding. Elife 4:e05345.

19. Kaufman J. 2018. Generalists and Specialists: A new view of how MHC class I molecules fight infectious pathogens. Trends Immunol 39:367–379.

20. Tregaskes CA, Kaufman J. 2021. Chickens as a simple system for scientific discovery: The example of the MHC. Mol Immunol 135:12–20.

21. Kaufman J, Völk H, Wallny HJ. 1995. A “minimal essential Mhc” and an “unrecognized Mhc”: two extremes in selection for polymorphism. Immunol Rev 143:63–88.

22. Koch M, Camp S, Collen T, Avila D, Salomonsen J, Wallny HJ, van Hateren A, Hunt L, Jacob JP, Johnston F, Marston DA, Shaw I, Dunbar PR, Cerundolo V, Jones EY, Kaufman J. 2007. Structures of an MHC class I molecule from B21 chickens illustrate promiscuous peptide binding. Immunity 27:885–99.

23. Tregaskes CA, Harrison M, Sowa AK, van Hateren A, Hunt LG, Vainio O, Kaufman J. 2016. Surface expression, peptide repertoire, and thermostability of chicken class I molecules correlate with peptide transporter specificity. Proc Natl Acad Sci U S A 113:692–7.

24. Li X, Zhang L, Liu Y, Ma L, Zhang N, Xia C. 2020. Structures of the MHC-I molecule BF2*1501 disclose the preferred presentation of an H5N1 virus-derived epitope. J Biol Chem 295:5292–5306.

25. Xiao J, Xiang W, Zhang Y, Peng W, Zhao M, Niu L, Chai Y, Qi J, Wang F, Qi P, Pan C, Han L, Wang M, Kaufman J, Gao GF, Liu WJ. 2018. An invariant arginine in common with MHC Class II allows extension at the C-Terminal end of peptides bound to chicken MHC class I. J Immunol 201:3084–3095.

26. Zhang J, Chen Y, Qi J, Gao F, Liu Y, Liu J, Zhou X, Kaufman J, Xia C, Gao GF. 2012. Narrow groove and restricted anchors of MHC class I molecule BF2*0401 plus peptide transporter restriction can explain disease susceptibility of B4 chickens. J Immunol 189:4478–87.

27. van Hateren A, Carter R, Bailey A, Kontouli N, Williams AP, Kaufman J, Elliott T. 2013. A mechanistic basis for the co-evolution of chicken tapasin and major histocompatibility complex class I (MHC I) proteins. J Biol Chem 288:32797–32808.

28. Walker BA, Hunt LG, Sowa AK, Skjødt K, Göbel TW, Lehner PJ, Kaufman J. 2011. The dominantly expressed class I molecule of the chicken MHC is explained by coevolution with the polymorphic peptide transporter (TAP) genes. Proc Natl Acad Sci U S A 108:8396–401.

29. Kaufman J. 2015. Co-evolution with chicken class I genes. Immunol Rev 267:56–71.

30. Kosmrlj A, Read EL, Qi Y, Allen TM, Altfeld M, Deeks SG, Pereyra F, Carrington M, Walker BD, Chakraborty AK. 2010. Effects of thymic selection of the T-cell repertoire on HLA class I-associated control of HIV infection. Nature 465:350–4.

31. Paul S, Weiskopf D, Angelo MA, Sidney J, Peters B, Sette A. 2013. HLA class I alleles are associated with peptide-binding repertoires of different size, affinity, and immunogenicity. J Immunol 191:5831–9.

32. Miura T, Brockman MA, Schneidewind A, Lobritz M, Pereyra F, Rathod A, Block BL, Brumme ZL, Brumme CJ, Baker B, Rothchild AC, Li B, Trocha A, Cutrell E, Frahm N, Brander C, Toth I, Arts EJ, Allen TM, Walker BD. 2009. HLA-B57/B*5801 human immunodeficiency virus type 1 elite controllers select for rare gag variants associated with reduced viral replication capacity and strong cytotoxic T-lymphocyte [corrected] recognition. J Virol 83:2743–55.

33. Schneidewind A, Brockman MA, Sidney J, Wang YE, Chen H, Suscovich TJ, Li B, Adam RI, Allgaier RL, Mothé BR, Kuntzen T, Oniangue-Ndza C, Trocha A, Yu XG, Brander C, Sette A, Walker BD, Allen TM. 2008. Structural and functional constraints limit options for cytotoxic T-lymphocyte escape in the immunodominant HLA-B27-restricted epitope in human immunodeficiency virus type 1 capsid. J Virol 82:5594–605.

34. Rizvi SM, Salam N, Geng J, Qi Y, Bream JH, Duggal P, Hussain SK, Martinson J, Wolinsky SM, Carrington M, Raghavan M. 2014. Distinct assembly profiles of HLA-B molecules. J Immunol 192:4967–76.

35. Bashirova AA, Viard M, Naranbhai V, Grifoni A, Garcia-Beltran W, Akdag M, Yuki Y, Gao X, O’HUigin C, Raghavan M, Wolinsky S, Bream JH, Duggal P, Martinson J, Michael NL, Kirk GD, Buchbinder SP, Haas D, Goedert JJ, Deeks SG, Fellay J, Walker B, Goulder P, Cresswell P, Elliott T, Sette A, Carlson J, Carrington M. 2020. HLA tapasin independence: broader peptide repertoire and HIV control. Proc Natl Acad Sci U S A 117:28232–28238.

36. Bacon LD. 1987. Influence of the major histocompatibility complex on disease resistance and productivity. Poult Sci 66:802–11.

37. Longenecker BM, Pazderka F, Gavora JS, Spencer JL, Ruth RF. 1976. Lymphoma induced by herpesvirus:resistance associated with a major histocompatibility gene. Immunogenetics 3:401–407.

38. Afrache H, Tregaskes CA, Kaufman J. 2020. A potential nomenclature for the Immuno Polymorphism Database (IPD) of chicken MHC genes: progress and problems. Immunogenetics 72:9–24.

39. Hosomichi K, Miller MM, Goto RM, Wang Y, Suzuki S, Kulski JK, Nishibori M, Inoko H, Hanzawa K, Shiina T. 2008. Contribution of mutation, recombination, and gene conversion to chicken MHC-B haplotype diversity. J Immunol 181:3393–9.

40. Kaufman J, Salomonsen J, Skjødt K. 1989. B-G cDNA clones have multiple small repeats and hybridize to both chicken MHC regions. Immunogenetics 30:440–51.

41. Kaufman J, Skjødt K, Salomonsen J. 1991. The B-G multigene family of the chicken major histocompatibility complex. Crit Rev Immunol 11:113–43.

42. Salomonsen J, Chattaway JA, Chan AC, Parker A, Huguet S, Marston DA, Rogers SL, Wu Z, Smith AL, Staines K, Butter C, Riegert P, Vainio O, Nielsen L, Kaspers B, Griffin DK, Yang F, Zoorob R, Guillemot F, Auffray C, Beck S, Skjødt K, Kaufman J. 2014. Sequence of a complete chicken BG haplotype shows dynamic expansion and contraction of two gene lineages with particular expression patterns. PLoS Genet 10:e1004417.

43. Gao C, Han L, Han J, Liu J, Jiang Q, Guo D, Qu L. 2015. Establishment of six homozygous MHC-B haplotype populations associated with susceptibility to Marek’s disease in Chinese specific pathogen-free BWEL chickens. Infect Genet Evol 29:15–25.

44. Shaw I, Powell TJ, Marston DA, Baker K, van Hateren A, Riegert P, Wiles MV, Milne S, Beck S, Kaufman J. 2007. Different evolutionary histories of the two classical class I genes BF1 and BF2 illustrate drift and selection within the stable MHC haplotypes of chickens. J Immunol 178:5744–52.

45. Hülsmeyer M, Fiorillo MT, Bettosini F, Sorrentino R, Saenger W, Ziegler A, Uchanska-Ziegler B. 2004. Dual, HLA-B27 subtype-dependent conformation of a self-peptide. J Exp Med 199:271–81.

46. Burrows SR, Rossjohn J, McCluskey J. 2006. Have we cut ourselves too short in mapping CTL epitopes? Trends Immunol 27:11–6.

47. Turner SJ, Kedzierska K, Komodromou H, La Gruta NL, Dunstone MA, Webb AI, Webby R, Walden H, Xie W, McCluskey J, Purcell AW, Rossjohn J, Doherty PC. 2005. Lack of prominent peptide-major histocompatibility complex features limits repertoire diversity in virus-specific CD8+ T cell populations. Nat Immunol 6:382–9.

48. Eldaghayes I, Rothwell L, Williams A, Withers D, Balu S, Davison F, Kaiser P. 2006. Infectious bursal disease virus: strains that differ in virulence differentially modulate the innate immune response to infection in the chicken bursa. Viral Immunol 19:83–91.

49. Kaiser P, Rothwell L, Galyov EE, Barrow PA, Burnside J, Wigley P. 2000. Differential cytokine expression in avian cells in response to invasion by Salmonella typhimurium, Salmonella enteritidis and Salmonella gallinarum. Microbiology (Reading) 146 Pt 12:3217–3226.

50. Kaiser P, Underwood G, Davison F. 2003. Differential cytokine responses following Marek’s disease virus infection of chickens differing in resistance to Marek’s disease. J Virol 77:762–8.

51. Liu WJ, Lan J, Liu K, Deng Y, Yao Y, Wu S, Chen H, Bao L, Zhang H, Zhao M, Wang Q, Han L, Chai Y, Qi J, Zhao J, Meng S, Qin C, Gao GF, Tan W. 2017. Protective T cell responses featured by concordant recognition of middle east respiratory syndrome coronavirus-derived CD8+ T cell epitopes and host MHC. J Immunol 198:873–882.

52. Otwinowski Z, Minor W. 1997. Processing of X-ray diffraction data collected in oscillation mode. Methods Enzymol 276:307–26.

53. Brünger AT, Adams PD, Clore GM, DeLano WL, Gros P, Grosse-Kunstleve RW, Jiang JS, Kuszewski J, Nilges M, Pannu NS, Read RJ, Rice LM, Simonson T, Warren GL. 1998. Crystallography & NMR system: A new software suite for macromolecular structure determination. Acta Crystallogr D Biol Crystallogr 54:905–21.

54. Emsley P, Lohkamp B, Scott WG, Cowtan K. 2010. Features and development of Coot. Acta Crystallogr D Biol Crystallogr 66:486–501.

55. Collaborative Computational Project, Number4, 1994. The CCP4 suite: programs for protein crystallography. Acta Crystallogr D Biol Crystallogr 50:760–3.

56. Adams PD, Afonine PV, Bunkóczi G, Chen VB, Davis IW, Echols N, Headd JJ, Hung LW, Kapral GJ, Grosse-Kunstleve RW, McCoy AJ, Moriarty NW, Oeffner R, Read RJ, Richardson DC, Richardson JS, Terwilliger TC, Zwart PH. 2010. PHENIX: a comprehensive Python-based system for macromolecular structure solution. Acta Crystallogr D Biol Crystallogr 66:213–21.

57. Laskowski RA, MacArthur MW, Moss DS, Thornton JM. 1993. PROCHECK: a program to check the stereochemical quality of protein structures. J Appl Crystallogr 26:283–291.

58. Thompson JD, Gibson TJ, Plewniak F, Jeanmougin F, Higgins DG. 1997. The CLUSTAL_X windows interface: flexible strategies for multiple sequence alignment aided by quality analysis tools. Nucleic Acids Res 25:4876–82.

59. Gouet P, Robert X, Courcelle E. 2003. ESPript/ENDscript: extracting and rendering sequence and 3D information from atomic structures of proteins. Nucleic Acids Res 31:3320–3.

